# FEDKEA: Enzyme function prediction with a large pretrained protein language model and distance-weighted k-nearest neighbor

**DOI:** 10.1101/2024.08.12.604109

**Authors:** Lei Zheng, Bowen Li, Siqi Xu, Junnan Chen, Guanxiang Liang

## Abstract

Recent advancements in sequencing technologies have led to the identification of a vast number of hypothetical proteins, surpassing current experimental capabilities for annotation. Enzymes, crucial for diverse biological functions, have garnered significant attention; however, accurately predicting enzyme EC numbers for proteins with unknown functions remains challenging. Here, we introduce FEDKEA, a novel computational method that integrates ESM-2 and distance-weighted KNN (k-nearest neighbor) to enhance enzyme function annotation. FEDKEA first employs a fine-tuned ESM-2 model with four fully connected layers to distinguish from other proteins. For predicting EC numbers, it adopts a hierarchical approach, utilizing distinct models and training strategies across the four EC number levels. Specifically, the classification of the first EC number level utilizes a fine-tuned ESM-2 model with three fully connected layers, while transfer learning with embeddings from this model supports the second and third-level tasks. The fourth-level classification employs a distance-weighted KNN model. Compared to existing tools such as CLEAN and ECRECer, two state-of-the-art computational methods, FEDKEA demonstrates superior performance. We anticipate that FEDKEA will significantly advance the prediction of enzyme functions for uncharacterized proteins, thereby impacting fields such as genomics, physiology and medicine. FEDKEA is easy to install and currently available at: https://github.com/Stevenleizheng/FEDKEA

## 1. Introduction

With the development of sequencing technologies, numerous hypothetical proteins are being discovered through ORF prediction tools(Hyatt, et al., 2010; Shendure, et al., 2017). The speed at which hypothetical proteins are discovered far exceeds the rate at which they can be experimentally annotated(Gill, et al., 2006; Qin, et al., 2010). For example, in 2023 alone, 29,526,946 protein sequences were uploaded to UniProt’s TrEMBL database (unreviewed), whereas only 1,699 sequences were added to the Swiss-Prot database (reviewed). Therefore, experimentally validated protein data represents only about 0.005% of predicted protein data(UniProt, 2021).

Enzymes, as one of the vital protein functions, have always been a focal point of research(Menendez-Arias, et al., 2017; Simpson, et al., 2024; Wang, et al., 2023). It is evident that experimentally characterizing enzyme functions of proteins is time-consuming and labor-intensive. Given the vast number of unannotated protein functions, there is an urgent need for new computational methods to annotate enzyme functions(Furnham, et al., 2009). Currently, enzyme function annotation for proteins is standardized using the Enzyme Commission (EC) number assigned by the International Union of Biochemistry and Molecular Biology Nomenclature Committee(IUBMBNC) (McDonald and Tipton, 2023). The IUBMBNC has classified over 6,800 enzymes, with a highly uneven distribution of data across enzyme classes. This disparity makes the accurate annotation of these enzymes’ EC numbers both a crucial and challenging task.

To address this issue, various computational methods have been developed for enzyme function annotation, including those based on sequence similarity(Altschul, et al., 1990; Desai, et al., 2011), homology modeling(Krogh, et al., 1994; Steinegger, et al., 2019), structure analysis(Zhang, et al., 2017), and machine learning(Ryu, et al., 2018; Sanderson, et al., 2023). The tool based sequence similarity, such as BLASTp, is widely used for protein function annotation by comparing unknown protein sequences with those annotated protein sequences. This similarity-based methods always cause low reliability when the sequence similarity is low. Moreover, sequence alignment approaches are often inadequate for capturing the intricate connections between protein structure and function. Machine learning models such as DeepEC and ProteInfer address enzyme function prediction by using multi-label classification and large-scale labeled datasets. However, the performance of these models is frequently hindered by poor generalization, limited accuracy, and insufficient coverage, primarily due to a lack of diverse and representative training data.

Transformer-based language models, initially developed for natural language processing (NLP), have been increasingly applied in the protein field to address various biological problems by treating protein sequences as a form of biological language(Elnaggar, et al., 2022; Lin, et al., 2023). Protein language model-based annotation shows unique advantages, effectively annotating low-similarity proteins with high throughput. These models employ pre-training on large protein datasets, gaining a comprehensive understanding of protein evolution, structure, and function. CLEAN and ECRECer are notable tools based on the protein language model ESM-1b for enzyme annotation(Shi, et al., 2023; Yu, et al., 2023). However, these two models have not been fine-tuned, resulting in their poor performance on enzyme annotation and limited applicability in practical scenarios.

Here, we propose a model that continues to utilize the protein language model-based strategy. Among various protein language models such as ESM and T5, the embeddings extracted by ESM, given the same parameter scale, have been found to be more conducive to downstream protein function classification(Thumuluri, et al., 2022). Recently, ESM released its second version, ESM-2, which outperforms ESM −1b in all aspects(Lin, et al., 2023; Rives, et al., 2021). Therefore, We fine-tune the ESM-2 model for specific tasks, including enzyme identification and EC number annotation, using a hierarchical classification strategy across the four levels of the EC numbering. The approach involves a series of models tailored for each level, employing transfer learning and a combination of MLP heads and distance-weighted KNN to ensure comprehensive enzyme annotation, even for classes with limited data. This approach aims to excel in EC number annotation without strictly relying on similarity.

## 2. Materials and methods

### 2.1 The dataset for model training

The dataset of UniProtKB SwissProt, released on March 2024, was collected for fine-tuning ESM-2 model and training MLP model. A total of 571,609 proteins was first filtered by sequence identity. A subset of 483,428 proteins containing 234,482 enzymes and 248,946 non-enzymes was split by the created time of proteins. For the binary classification task of determining whether a protein is an enzyme, protein data up to 2024 will be divided into training, validation, and test sets in an 8:1:1 ratio. Protein data from after 2024 will be used as an independent test set to evaluate the model’s performance. For the EC number classification tasks at various levels, we will use 234,482 enzyme sequences for model training and subsequently remove multi-functional enzymes. For the classification tasks of level 1 and level 2 EC numbers, we will similarly split the enzyme data up to 2024 into training, validation, and test sets in an 8:1:1 ratio, and use post-2024 enzyme data as an independent test set to assess the performance of the models.

Due to the scarcity of some enzyme classes, we will not create an independent test set based on the year for level 3 and level 4 EC number classification tasks. Instead, we will split the data into training and validation sets in an 8:2 ratio to determine the optimal K value for the KNN model.

### 2.2 Model structures and training processes

The overall framework of the model involves fine-tuned ESM-2 and distance-based KNN enabled enzyme annotation (FEDKEA). The structures of FEDKEA are shown in **Fig.3**. The model framework consists of two main parts: determining whether a protein is an enzyme and predicting the enzyme’s EC number. For the binary classification task of determining if a protein is an enzyme, we use the ESM-2 model with 33 layers and 650M parameters. First, the amino acid sequence of the protein is tokenized. We then fine-tune the weights of the last few layers, finding that fine-tuning four layers yields the best performance. The embeddings from the fine-tuned model are averaged according to the sequence length, resulting in a 1280-dimensional vector. This vector is fed into a five-layer MLP (1280-960-480-120-30-2), with each layer using ReLU activation for further feature extraction. The output of the MLP is then passed through a softmax layer to calculate the probability for each class, classifying a protein as an enzyme if the probability is ≥0.5 and not an enzyme otherwise.

For the EC number classification task, we adopt a hierarchical prediction strategy based on the four-level structure of EC numbers. For the first-level classification, a seven-class task, we use the same strategy as the binary classification: fine-tuning the 33-layer, 650M parameter ESM-2 model, and find that fine-tuning three layers yields the best performance. The MLP for this task is adjusted to four layers (1280-960-480-120-classes), and considering the class imbalance, we add a batch normalization layer after the MLP, followed by ReLU activation. The binary cross-entropy loss function is replaced with Focal Loss to solve the problem of class imbalance.

For the second and third-level EC number classification tasks, we use the same model framework, incorporating transfer learning to utilize embeddings learned from the first-level classification. Additionally, enzyme classes with few samples are grouped into an “others” category. For the fourth-level classification, due to the scarcity of enzymes in most classes, we use a distance-weighted KNN model, inheriting the embeddings from the first-level classification. Testing revealed that K=3 yielded the best performance.

Throughout the training process, we employ early stopping and the Adam optimizer (Kingma and Ba, 2015), with a learning rate of 5e-6 and a weight decay of 1e-5.

### 2.3 Test process and Evaluation metrics

For the binary classification task of determining if a protein is an enzyme, the accuracy (ACC), precision, recall, F1, AUC, and AP value. The formulas for ACC, precision, recall, and F1 are as follows (1)-(4):

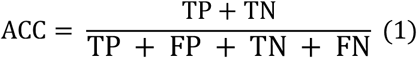

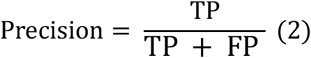

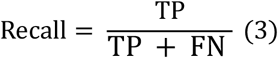

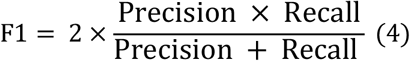

where TP, FP, TN, and FN mean the number of true positive, false positive, true negative, and false negative samples during a test. The metrics mentioned above are typically calculated assuming a probability threshold of 0.5. However, altering the classification threshold results in different metric values. By evaluating these metrics at various thresholds, ROC and PR curves can be plotted. The ROC curve illustrates the relationship between the true positive rate (TPR) and false positive rate (FPR), while the PR curve demonstrates the relationship between precision and recall. The formulas for TPR and FPR are given in equations (5) and (6).

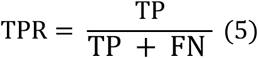

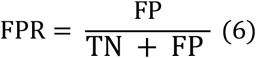

The AUC represents the area under the ROC curve, and the AP represents the area under the PR curve. Combining the above metrics could evaluate and analyze the performance of different models from multiple perspectives, especially the F1, AUC and AP metrics.

For multiple classification problems, the evaluation criteria included mACC (macro-average accuracy), mPR (macro-average precision), mRecall (macro-average recall), and mF1(macro-average F1 value). These formulas are given in equations (7) - (10).

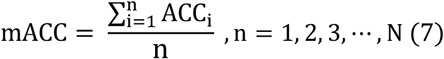

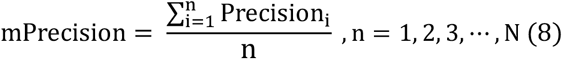

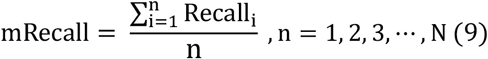

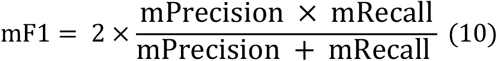

We will use all protein data uploaded in 2024, totaling 415 proteins, including 148 enzymes and 267 non-enzyme proteins, as an independent test set to validate the model’s accuracy. Additionally, when validating the EC number of enzymes, considering the presence of multi-functional enzymes, we define a prediction as correct if any one of the enzyme’s functions is correctly predicted.

### 2.4 Computing resources

Up to eight 40G NVIDIA A40 and two 32G NVIDIA Tesla V100 PCle GPUs were utilized for model training and inference, and these GPUs were all from the public platform of School of Medicine, Tsinghua University.

## 3. Results

### 3.1 Model development and evaluation

The overall framework of the model involves fine-tuned ESM-2 and distance-based KNN enabled enzyme annotation (FEDKEA). Initially, protein sequences are subjected to analysis within the fine-tuned ESM-2 model, where the last four layers are specifically adapted to discern enzymatic attributes. Following this initial assessment, should the protein be identified as an enzyme, it undergoes further embedding step within the ESM-2 model, wherein the last three layers are fine-tuned. During this process, embedding data are shared globally, and subsequently subjected to diverse Multi-Layer Perceptron (MLP) models. Ultimately, the processed data are fed into a KNN model, utilizing distance weighting, to ascertain the final prediction at the concluding stage (**Fig 1**).

**Fig. 1.**
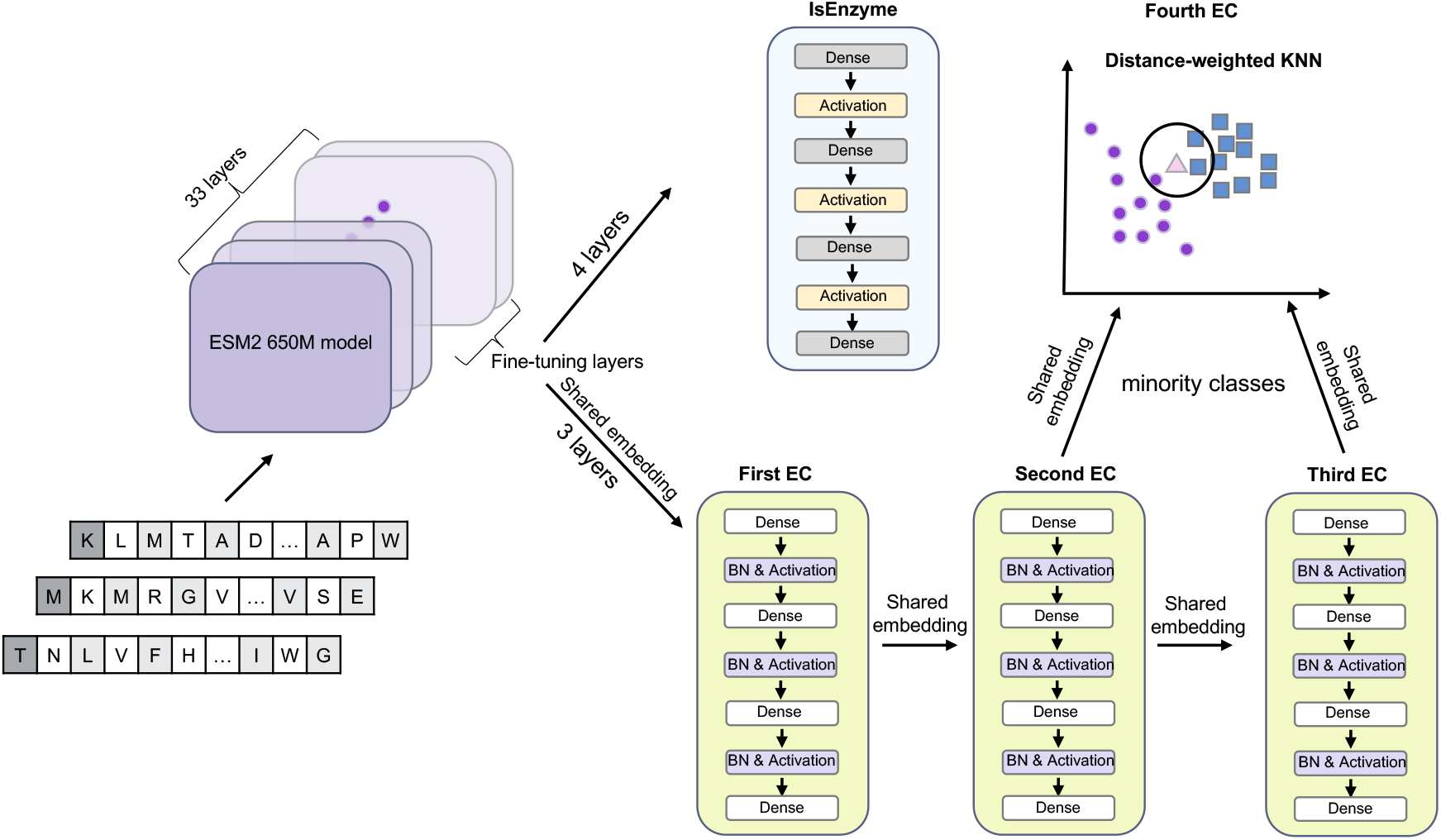
The fine-tuned ESM-2 and distance-based KNN framework of FEDKEA. The model employs fine-tuned ESM-2 for enzyme detection, followed by global sharing of embedding data and MLP processing, culminating in KNN-based prediction.

With the development of the large language model (LLM), ESM-2 model as a state-of-the-art protein language model, at scales from 8 million parameters up to 15 billion parameters, is trained to predict the identity of amino acids that have been randomly masked out of protein sequences. ESM-2 model is used to help us acquire rich protein information embedded inside the protein sequence. Since our task is enzyme commission annotation rather than protein structure prediction, we used a fine-tuned ESM-2 model trained by annotated proteins with enzyme function so that it could help us extract more protein information about enzyme functions. Considering the hierarchical organization of enzyme classification, which spans four levels with increasing specificity and a sparse distribution of categories at the final level, we have adopted a hierarchical approach. By integrating this strategy with distance-weighted K-nearest neighbor (KNN) algorithms, our goal is to enhance the accuracy of enzyme function prediction, particularly at the fourth level.

In the training stage, a universal protein knowledgebase UniProt released before 2024 was used for model development and evaluation. For both the enzyme identification and enzyme commission first-level classification tasks, we fine-tuned the ESM-2 using data before 2021 and evaluated its performance using a validation set from 2021-2023, achieving a best 91.27% F1 score under fine-tuned last four layers of ESM-2 (**Fig 2A, 2B**) and a best 88.27% F1 score under fine-tuned last three layers of ESM-2 (**Fig 3A, 3B, 3C**), respectively. The layer 33 as the embedding data better improves the ability of model enzyme identification than the layer 32 as the embedding data (**Fig 2C**).

**Fig. 2.**
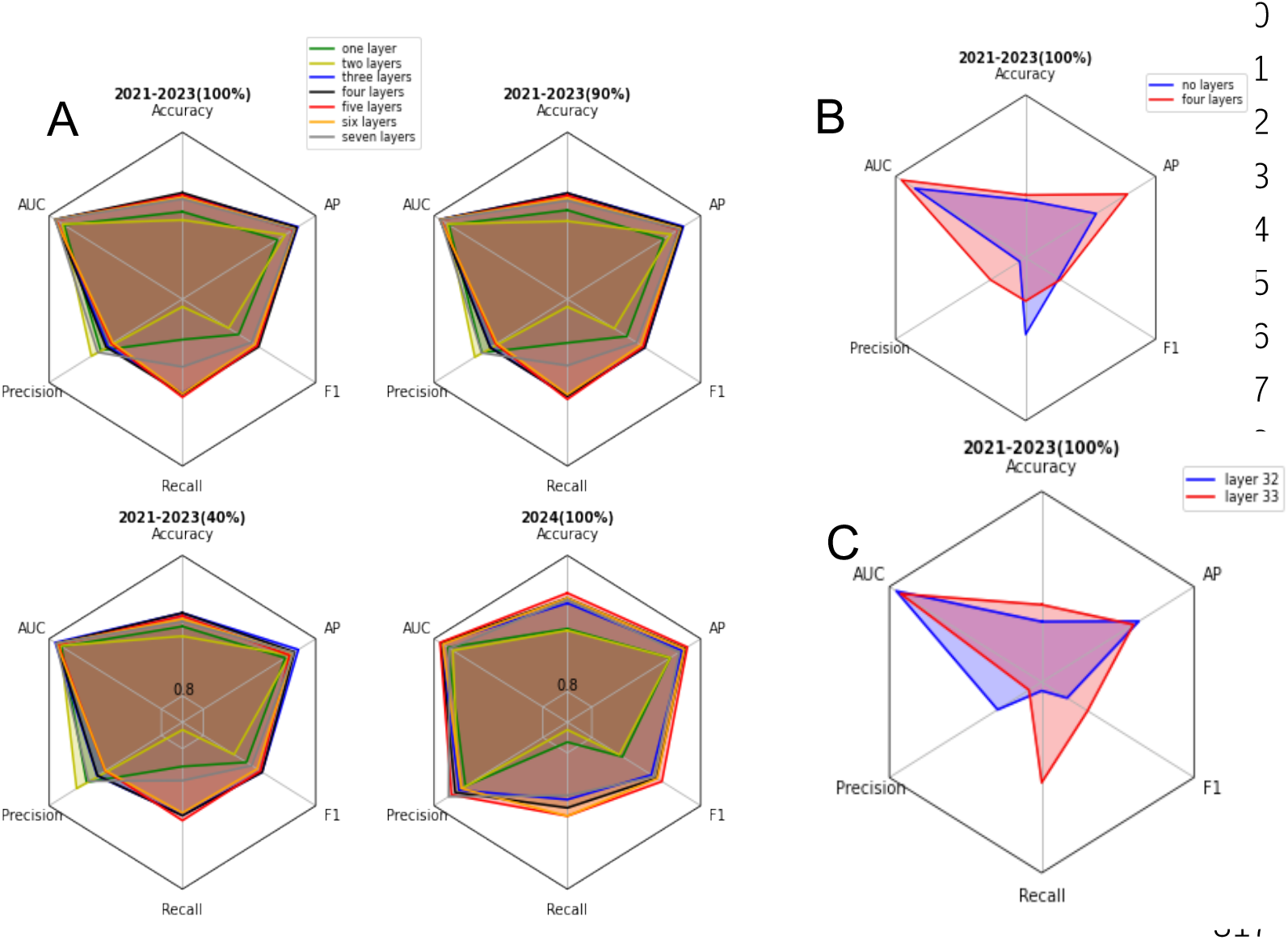
Fine-tuned model for the enzyme identification. (A) The performance metrics (accuracy, precision, recall, AUC, AP, and F1 score) of the fine-tuned model on datasets from 2021-2023 and 2024 are evaluated across layers 1-7. (B) The performance metrics (accuracy, precision, recall, AUC, AP, and F1 score) comparison of model between fine-tuned four layers and fine-tuned zero layer on datasets from 2021-2023. (C) The performance metrics (accuracy, precision, recall, AUC, AP, and F1 score) comparison of models with the 32nd layer as the embedding layer and the 33rd layer as the embedding layer on datasets from 2021-2023

**Fig. 3.**
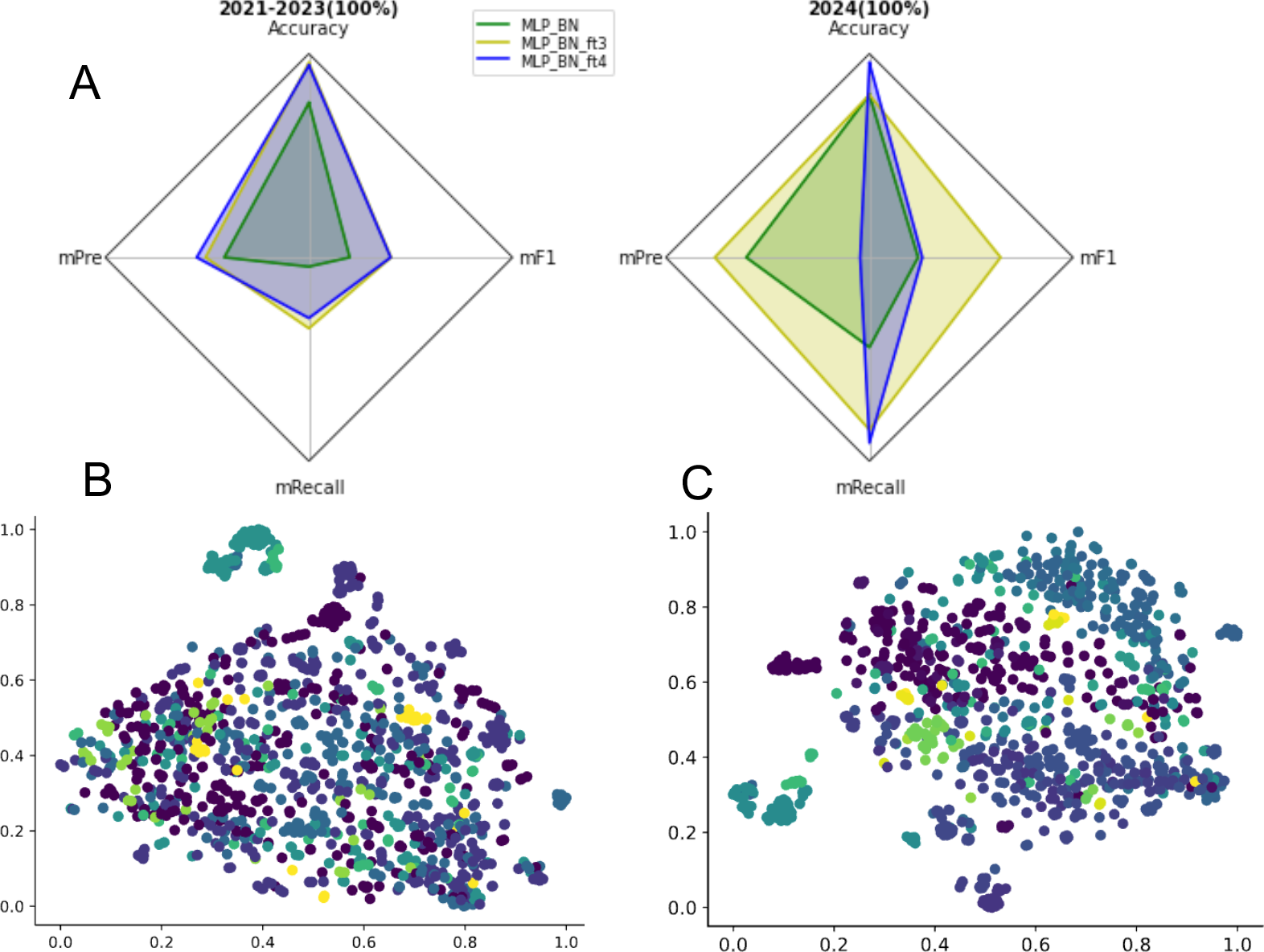
Fine-tuned model for enzyme commission first-level classification. (A) The performance metrics (accuracy, mPrecision, mRecall, and mF1 score) of the fine-tuned model on datasets from 2021-2023 and 2024 are evaluated across layers 3-4. (B) The tSNE plot of the 33rd layer in the non-fine-tuned model. (C) The tSNE plot of the 33rd layer in the model fine-tuned on four layers.

### 3.2 Benchmarking FEDKEA with previous EC number annotation tools

After training, the prediction performance of FEDKEA was systematically investigated by comparing it with two recently published state-of-the-art deep learning-based EC number annotation tools [i.e., CLEAN and ECRECer]. One independent dataset, named UniProtKB_2024_02, consisted of 172 enzyme sequences and 243 non enzyme sequences that not included in any model’s development. The prediction scenario fully represented a practical situation, where the labeled knowledgebase was the Swiss-Prot database and related enzyme information of query sequences were unknown. In the enzyme identification task, FEDKEA achieved the highest value in various multilabel accuracy metrics, including accuracy (0.9205), precision (0.9542) and F1 (0.8985) (**Fig 4A**). It is worth noting that CLEAN is not capable of recognizing enzymes.

**Fig. 4.**
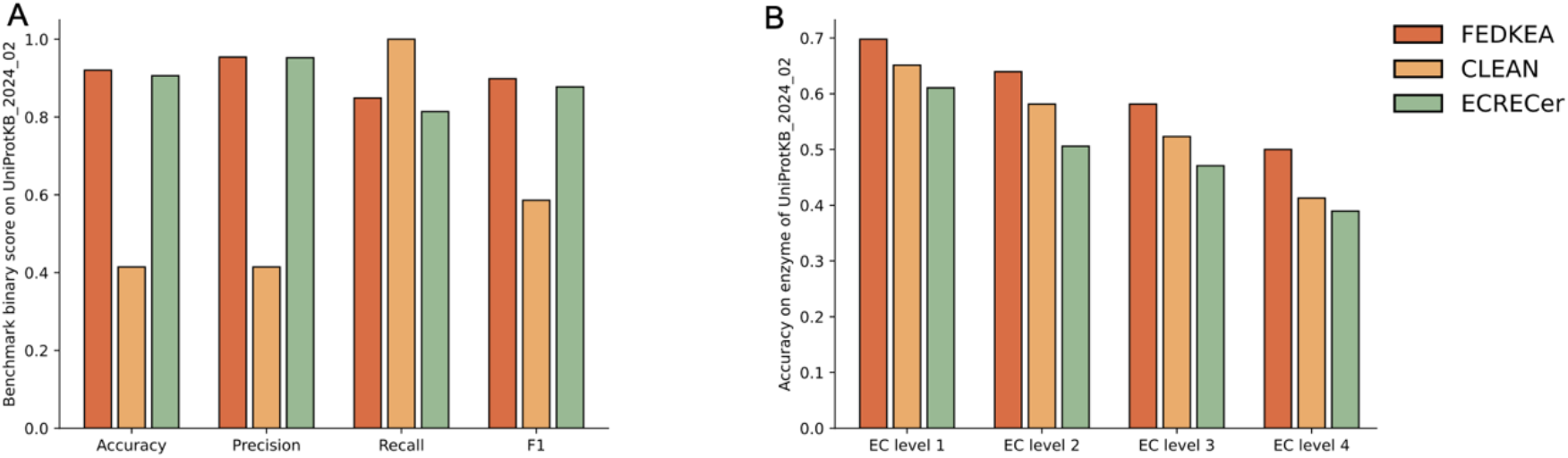
Quantitative comparison of FEDKEA with the state-of-the-art EC number prediction tools. (A) Evaluation of FEDKEA’s performance toward four multilabel accuracy metrics (accuracy, precision, recall and F1 score) in the task of enzyme identification on the UniProtKB_2024_02 dataset. (B) Accuracy comparison of FEDKEA, CLEAN and ECRECer at each level of enzyme commission number on the UniProtKB_2024_02 dataset.

We then compared the predictive performance of three tools at each level of enzyme commission number under the assumption that the protein sequence is an enzyme (172 enzyme sequences). Overall, FEDKEA resulted in better prediction accuracy (69.77 to 50%) compared with CLEAN (65.12 to 41.28%) and ECRECer (61.05 to 38.95%) (**Fig 4B**).

## 4. Discussion

Protein function annotation has long been a challenging problem in biology. Enzymes, as proteins involved in various biological processes, have consistently attracted the attention of researchers. Accurate annotation of enzyme functions remains a significant challenge. While experimental methods can precisely annotate enzyme functions, they are time-consuming and labor-intensive. Enzyme function annotation methods based on sequence similarity, homology, and structural alignment have been employed but often fall short in accurately predicting enzyme functions, particularly specific EC numbers.

With the rise of large language models in the past five years, there is new potential to further understand the rich information embedded within protein sequences from an evolutionary perspective. Our model addresses this by incorporating a protein large language model in its first module, using a fine-tuning strategy to tailor it for specific enzyme function annotation tasks. Additionally, recognizing the hierarchical nature of enzyme numbering, we have implemented a tiered approach to maintain high accuracy at each level of prediction. For the fourth-level categories, where data distribution is imbalanced, we use a distance-weighted K-nearest neighbor (KNN) model for final classification. This design enables our model to outperform other protein large language model-based tools, such as CLEAN and ECREC, in terms of generalization and prediction accuracy on unknown datasets. However, during the model training process, we remain reliant on well-annotated enzyme data, and the model still struggles to provide high-confidence results for novel proteins. Additionally, almost models, including our model, only consider the prediction of enzyme function but not predict the catalytic site of enzyme.

